# Mitochondrial genetic effects on reproductive success: signatures of positive intra-sexual, but negative inter-sexual pleiotropy

**DOI:** 10.1101/138180

**Authors:** M. Florencia Camus, Damian K. Dowling

**Affiliations:** School of Biological Sciences, Monash University, 3800, Australia; Research Department of Genetics, Evolution and Environment, University College London, Gower Street, London, WC1E 6BT, United Kingdom

**Keywords:** mitochondria, life-history, sexual conflict, reproduction, maternal inheritance, sexual antagonism

## Abstract

Mitochondria contain their own DNA, and numerous studies have reported that genetic variation in this (mt)DNA sequence modifies the expression of life-history phenotypes. Maternal inheritance of mitochondria adds a layer of complexity to trajectories of mtDNA evolution, because theory predicts the accumulation of mtDNA mutations that are male-biased in effect. While it is clear that mitochondrial genomes routinely harbor genetic variation that affects components of reproductive performance, the extent to which this variation is sex-biased, or even sex-specific in effect, remains elusive. This is because nearly all previous studies have failed to examine mitochondrial genetic effects on both male and female reproductive performance within the one-and-the-same study. Here, we show that variation across naturally-occurring mitochondrial haplotypes affects components of reproductive success in both sexes, in *Drosophila melanogaster*. However, while we uncovered evidence for positive pleiotropy, across haplotypes, in effects on separate components of reproductive success when measured within the same sex, such patterns were not evident across sexes. Rather, we found a pattern of sexual antagonism across haplotypes on some reproductive parameters. This suggests the pool of polymorphisms that delineate global mtDNA haplotypes is likely to have been partly shaped by maternal transmission of mtDNA and its evolutionary consequences.

## Introduction

Eukaryotic cells are thought to have arisen from the ancient symbiotic union between two prokaryote cells; one an α-proteobacterium and the other an archean-like organism. The α-proteobacterium would evolve into the mitochondrion, and the archaea-like bacterium into the eukaryote cell (1). Moreover, each of these ancestral entities possessed their own genomes, and their symbiosis kick-started millions of years of inter-genomic coevolution that delineates contemporary eukaryotes from the organisms of other domains (2). Almost without exception, eukaryotes from fungi to animals have retained these two genomes – one mitochondrial (comprised of mtDNA), the other nuclear, and interactions between genes spanning each of these genomes coordinate closely to regulate critical biological processes tied to cellular metabolism via oxidative phosphorylation (OXPHOS) (3-5).

Over the course of evolutionary history, most of the genes in the mitochondrial genome have translocated to the host nuclear genome, leaving only a small number of genes in the mtDNA, including thirteen protein-coding genes that are salient to OXPHOS function (4). Evolutionary biologists long assumed that purifying selection would prevent any non-neutral (i.e., phenotype-modifying) genetic variation from accumulating within these mtDNA-encoded genes, given these genes encode essential subunits of the electron transport chain, and given that the mitochondrial genome is haploid and therefore all alleles within it are invariably exposed to natural selection (6-8). As such, the mitochondrial genome was harnessed as the quintessential molecular marker upon which to base evolutionary and population genetic inferences, facilitated by its high mutation rate, maternal inheritance and general lack of recombination (6, 9-12).

Over the past two decades, however, an increasing number of studies has challenged this assumption of neutrality (7, 13, 14). In particular, numerous studies have used multigenerational breeding schemes with the power to partition mitochondrial genetic from nuclear genetic effects, and revealed that the genetic polymorphisms that delineate distinct mitochondrial haplotypes, sourced from separate populations, contribute to the expression of life-history traits tied to reproductive success, development, and longevity (3, 15-23).

Currently, however, it is unclear how these phenotype-modifying genetic polymorphisms accumulate within mitochondrial genomes. One alternative is that they constitute adaptations, fixed under natural selection. This is consistent with the results of some studies that examined mutational profiles of mtDNA sequences, and found signatures of positive selection in the form of elevated ratios of non-synonymous (*dN* – changes amino acid) to synonymous (*dS* – does not change amino acid) mutations (24-26). Alternatively, such polymorphisms might rise to appreciable frequencies within populations under mutation-selection balance, and potentially then be fixed by drift. This alternative is plausible; firstly, given the mitochondrial genome has a high mutation rate relative to its nuclear counterpart (27). Secondly, it is reasonable to predict there will be a diminished efficiency of selection in shaping the mtDNA sequence relative to nuclear DNA sequences, because of a theorised fourfold reduction in the effective population size of the mitochondrial genome that stems from it being haploid and maternally inherited (6, 7).

Maternal inheritance of mitochondrial genomes adds a further layer of complexity to the dynamics of mtDNA sequence evolution, because it means that selection can only act on non-neutral mtDNA polymorphisms directly through the female lineage (28-30). This hypothesis, which has been called “Mothers Curse” (29), predicts that mutations that are neutral, beneficial or even slightly deleterious to females may accumulate in the mtDNA sequence even if these very same mutations are harmful in their effects on males (30). Recent studies in *Drosophila* uncovered evidence for the existence of a pool of male-harming, but female-neutral polymorphisms that have accrued within mtDNA haplotypes, and which affects genome-wide patterns of gene expression in males, particularly of genes involved in encoding male-specific reproductive tissues (31), and that shapes patterns of male, but not female, longevity (21, 32).

However, the extent to which mitochondrial haplotypes exhibit sex-biases in their effects on the expression of life history phenotypes remains unclear, because relative few studies have measured phenotypic effects associated across sets of naturally-occurring mtDNA genotypes in both males and females, respectively (18, 21-23, 31-35). The sparsity of studies reporting sex-specificity in effects is particularly evident when it comes to traits tied to reproductive performance. Indeed, we are aware of only a single study to date that sought to measure mtDNA-mediated effects on components of reproductive success in both males and females. Immonen et al. (2016) examined the expression of components tied to reproductive success in each of the sexes across orthogonal combinations of mitochondrial and nuclear genotype sourced from three distinct populations, in the seed beetle, *Callosobruchus maculatus*. The nuclear genomic backgrounds, against which the three different mtDNA haplotypes were placed, were not isogenic, but rather represented by large pools of segregating nuclear allelic variance that were sourced from each of three global populations. The authors reported mitochondrial genetic, and mito-nuclear interactions for female fecundity, and male ejaculate weight, and also an effect on female egg size that was traceable to an interaction involving the age and mito-nuclear genotype of the sire. Correlations in the reported mitochondrial, or mito-nuclear, genetic effects across the measured traits were, however, not examined (22)

Broadly, the general failure of previous studies to have examined mitochondrial genetic contributions to reproductive phenotypes in both sexes has led to a gap in our understanding of the genetic architecture of mitochondrial genomes, particularly in light of the prediction that traits and tissue types exhibiting strong sexual dimorphism and sex-limitation in expression (such as the testes, sperm and reproductive glands involved in male reproductive outcomes) are hypothesized to be the key candidates for susceptibility to Mother's Curse effects (28, 30, 31, 36). Furthermore, very few studies have measured multiple traits across the same set of mtDNA genotypes, to examine levels and patterns of mtDNA-linked pleiotropy across traits, within and across the sexes. Studies that have screened for such pleiotropy have, however, reported interesting patterns, providing insights into the evolutionary processes by which genetic variation can accumulate within mitochondrial genomes. For example, Dowling et al. (2009) found a strong positive association in effects of two mtDNA haplotypes segregating within a population of *D. melanogaster*, on two life history traits in females - reproductive performance and longevity (37). The haplotype conferring higher female reproductive success also conferred higher female lifespan. In another study, Camus et al (2015) reported that a SNP found within the mtDNA-encoded *CYTB* gene of *D. melanogaster*, which is associated with low fertility in males (38) but not in females, confers higher male lifespan but shorter female lifespan relative to haplotypes harbouring other variants of this gene (21). This SNP is therefore associated with antagonistic pleiotropic effects both within and across the sexes, consistent with the idea that some mtDNA SNPs might accumulate under positive selection in females, even if they are associated with suboptimal male phenotypes (30). If so, then maternal inheritance of mitochondria could potentially lead to sexually antagonistic trajectories of mtDNA evolution (39-41).

To address patterns of sex-specificity and pleiotropy of mitochondrial genetic variation, here we screen thirteen naturally-occurring mitochondrial haplotypes of *D. melanogaster*, each sourced from a distinct global population, for components of reproductive output in each sex. We used strains in which each of these haplotypes had been placed alongside an isogenic nuclear background prior to the phenotypic assays (32, 42, 43), such that all phenotypic effects observed could be traced directly to genetic polymorphisms separating each haplotype. Firstly, we measured reproductive success of males and females who had abstained from sexual interactions until the peak of their fertility, and were then provided with a 24 h opportunity to mate and reproduce (hereafter termed “short-burst” components of reproduction). Secondly, we measured reproductive success of each sex over a prolonged period of time, from eclosion into adulthood to 8 (male) and 12 (female) days of age (termed “sustained” reproductive success). Thus, we performed two assays of reproductive success for each sex – one representing success based on a limited opportunity at the peak of an individual's reproductive lifespan; the other based on reproductive stamina when faced with multiple opportunities and partners across the early phase of adult life.

## Materials and Methods

### Mitochondrial lines

Thirteen *Drosophila melanogaster* strains were used, and these strains have been previously described (38, 43). In brief, the isogenic nuclear background from the *w^1118^* strain (Bloomington stock number: 5905) was coupled to mitochondrial haplotypes from thirteen distinct geographic locations using a crossing scheme that is outlined in Clancy (2008). These strains have each been maintained in duplicate since 2007, with the duplicates propagated independently, to enable us to partition mitochondrial genetic effects from cryptic nuclear variance that might have accumulated among the strains, as well as from other sources of environmental variation. Each generation, virgin females are collected from each duplicate of each mitochondrial strain (hereafter *mitochondrial strain duplicate*) and backcrossed to males of the *w^1118^* strain, to maintain isogenicity of the nuclear background. Furthermore, *w^1118^* is itself propagated by one pair of full-siblings per generation. Thus, if mutations arise in the *w^1118^* strain, they will be swiftly fixed and passed to all mitochondrial strain duplicates, thus maintaining the critical requirement of isogenicity of the nuclear genome.

One of the mitochondrial strains (Brownsville) incurs complete male sterility in the *w^1118^* nuclear background, whereas females who harbour this haplotype remain fertile (38). This strain was therefore excluded from assays of male reproductive success (n=12 haplotypes in these assays), but included in assays of female reproductive success (n=13 haplotypes). All mitochondrial strains and *w^1118^* flies were reared at 25°C, under a 12h: 12h light: dark photoperiod regime, on potato-dextrose-agar food medium and with *ad libitum* access to live yeast. All strains had been cleared of any potential endosymbionts, such as *Wolbachia*, through tetracycline treatment at the time that the strains were created (44). Diagnostic PCR with *Wolbachia*-specific primers confirmed all lines are free of *Wolbachia* (45).

### Male Reproductive Success

Two separate components of male reproductive success were measured, via two separate experiments. The first experiment measured male short-burst offspring production, i.e. following an exposure to a single female at the peak age of male reproductive fertility. This assay measures the ability of a male to convince a virgin female to mate, and then measures the number of offspring produced from sexual interaction with that female, which is likely to be a function of the males ejaculate quality (number and quality of sperm, and content and quality of reproductive proteins, transferred). The second experiment gauged sustained offspring production across the first eight days of adult life, during which time males had ongoing access to new and virgin females. This assay thus represents a measure of male reproductive stamina (a function of male mating rate across time, and ability to replenish sperm and ejaculate stores). Each assay is described below.

#### Male reproductive success following exposure to a single female (short-burst offspring production)

This experiment measured offspring produced by a single male after a one-off mating opportunity with a virgin female when 4 days of adult age. The assay was run in two blocks, each separated in time by one generation. For three generations leading up to the experiment, each mitochondrial strain duplicate was propagated across 3 vials, with each vial containing 10 pairs of flies of standardised age (4 day old), and at controlled larval densities (approximately 80 eggs per vial). Then, ten virgin males from each mitochondrial strain duplicate (total 20 male flies per haplotype) were collected randomly from the 3 vials that propagate the line, and each stored individually in separate 40 ml vials containing 5mL of food medium. At the same time, virgin females were collected from the isogenic *w^1118^* strain to be used as “tester” flies in the experiment. These females were sourced from 10 separate vials, which had been propagated and stored under the same experimental conditions as described for the mitochondrial strain focal males, and they were stored in groups of 10 females per vial.

When four days old, each focal male was then combined with an equivalently-aged “tester” female, and these flies then cohabited the same vial for a 24 h period. Following this, focal males were removed from the mating vial and discarded. Females were then transferred into fresh vials with food substrate every 24 h over a 4 d period. The total number of offspring eclosing across these four vials was recorded for each focal male.

#### Male reproductive across 8 days (sustained offspring production)

Sustained offspring production was assayed following the method described in Yee et al. (2015). In brief, individual males collected from each mitochondrial strain duplicate were provided with the opportunity to mate with eight different virgin females over eight consecutive 24 h long exposures (46). To initiate the assay, twenty virgin males were collected from each mitochondrial strain duplicate, and each placed in a separate vial (total of 40 flies per mitochondrial haplotype). Twenty-four hours later, one 4-day-old virgin *w^1118^* female was added to each vial, and the focal male and tester female then cohabited for 24 h. Following this 24 h exposure, males were removed and placed with another 4-day-old virgin w*^1118^* female for another 24 h period. This process was repeated until day eight of the experiment (8 separate exposures). After each exposure, the *w^1118^* females were retained and themselves transferred into fresh vials every 24 h for a total period of 4 consecutive days (including the 24 h cohabitation period), thus providing each female with up to 96 h to oviposit. Thirteen days following the 96h oviposition period, the number of eclosed adult offspring emerging from each vial was counted.

### Female reproductive success

Two separate components of female reproductive success were measured. The first experiment gauged “short-burst” components of success, in which the number of eggs produced per female (fecundity), number of adults (reproductive success) produced, and proportion of eggs that ultimately eclosed into adulthood (an index of short-burst viability) were scored, following a 24 h laying opportunity at the peak age of female fecundity (4 days of age). The second experiment measured sustained performance, in which female reproductive success was calculated over a 13-day period, thus representing a measure of reproductive stamina.

#### Female components of short-burst offspring production, and short-burst ‘egg-to-adult’ viability

The assay was run in five blocks, each separated in time by one generation. Female focal flies from each mitochondrial strain duplicate were collected as virgins, and stored individually. These were collected over numerous 40mL vials, each of which had been propagated by 10 pairs of age-controlled parents (4 day old), and at controlled larval densities (approximately 80 eggs per vial). When 4 days of age, each female was exposed to one 4 d old tester virgin male, collected from the *w^1118^* strain, for a period of 12 hours and then the females transferred to a fresh vial for 24 h to oviposit. Following this 24 hour ovipositioning period, females were discarded. We counted the eggs oviposited per female over this 24 h period (an index of short-burst fecundity), plus the offspring that emerged from these eggs (an index of short-burst offspring production).

Furthermore, we were able to calculate the proportion of eggs laid by each female that were converted into adult offspring (short-burst viability). Although many studies treat egg-adult viability as a measure of offspring viability, this trait stands at the nexus between a maternal and an offspring trait (47), and it is well established that maternal effects affect the trait in *D. melanogaster* (48-50). Indeed, strong maternal effects on this trait are in direct alignment with predictions of classic life-history theory, in which maternal resource provisioning into the ova lies at the heart of the classic evolutionary trade-off between gamete size and number (51); a trade-off that extends to *Drosophila* (52, 53). While ultimately it is not possible for us to delineate whether any mitochondrial haplotype effects on short-burst viability are manifested primarily through mothers (as mtDNA-mediated maternal effects) or primarily on the offspring themselves (via the direct effects of mtDNA mutations on survival through juvenile development), it is nonetheless highly informative to examine patterns of mitochondrial haplotypic variation affecting this trait among the other dedicated components of female and male reproductive success, and thus we include it in our study.

#### Female offspring production across 13 days (sustained offspring production)

Forty females from each mitochondrial strain duplicate were collected as virgins, and placed in individual vials. One day later, two 4 d old virgin *w^1118^* males were placed into each female vial. Females, and the two males with which each female cohabited, were then transferred into fresh vials every 24 hours, for 13 days. The accompanying males were discarded every fourth day, and two 4 d old virgin males of the *w^1118^* strain were added. This ensured that females were not sperm-limited throughout the duration of the experiment. At the end of day 13, all flies across all vial were discarded, and vials were kept for eggs to develop. Female reproductive success was determined by counting the total number of adult offspring produced by each female, per vial, over the 13-day assay.

### Statistical Analysis

General linear mixed models, using a Gaussian distribution, were fitted to the male and female short-burst offspring production data. Female short-burst fecundity data was modelled by fitting a generalized linear mixed model, using a Poisson distribution. For data that conformed to a Poisson distribution, we checked for over-dispersion using the function “*dispersion_glmer*” in the package *blmeco* (54). Short burst viability data was modelled as a binomial vector, composed of the number of adults and number of eggs that failed to hatch (eggs-adults), using a binomial distribution and logit link. For each analysis, mitochondrial strain was modelled as a fixed effect, and the duplicate nested within mitochondrial strain and the sampling block (for assays of short-burst components, which were assayed over multiple blocks) included as random effects, in the *lme4* package (55) in R (56). The fitted models were evaluated by type III (Wald chisquare tests) sums of squares analysis of variance using the *car* package in R. We confirmed significance of individual factors using model comparison. The fitted models were evaluated by simplifying a full model, by sequentially removing terms that did not change the deviance of the model (at α = 0.05); starting with the highest order interactions, and using log-likelihood ratio tests to assess the change in deviance in the reduced model relative to the previous model (56).

For the experiments gauging sustained offspring production, the overall total number of offspring (for both male and female models) was zero-inflated, and the resulting models over-dispersed. We therefore analysed both datasets using a negative binomial distribution (57), in which the zero values are a blend of sampling and structural effects (negative binomial parameter; variance = ϕµ). These models were performed using the R (v. 3.0.2) package glmmADMB (http://glmmadmb.r-forge.r-project.org/glmmADMB.html). The response variable was total number of offspring produced, with mitochondrial strain and day of sampling, plus their interaction, as fixed factors. The random effect in the model was mitochondrial duplicate nested within mitochondrial strain.

A matrix of mitochondrial genetic correlations (Pearson's correlation coefficients) was created by obtaining mtDNA haplotype-specific means for each reproductive trait across all mitochondrial strains. Thus, we had 13 means (one per haplotype) for each female measure of short burst (including short-burst viability) and sustained offspring production, and 12 means for the male measures (since the Brownsville haplotype was excluded from the male assays). Inter-sexual correlations across haplotypes were thus based on 12 means. Correlation coefficients of all pairwise combinations of traits were then evaluated using a bootstrapping procedure, in which trait means were resampled with replacement (1000 replicates), and 95% confidence intervals were calculated using the Adjusted Percentile (BCa) Confidence interval method. Bootstrapped correlation coefficients plus their confidence intervals were calculated using the functions “*boot”* and “*boot.ci”* in the R package *boot* (58). Correlations with confidence intervals that did not overlap with zero were considered statistically significant.

## Results

### Male Mitochondrial Reproductive Success Assays

The identity of the mitochondrial strain affected male short-burst offspring production (χ^2^ = 30.992, p = 0.001, Table 1A). Male sustained offspring production was affected by an interaction between mitochondrial strain and day of mating (haplotype × day, χ^2^ = 183.039, p<0.001, Table 1B, Figure 1A, Figure 1A). Male offspring production tended to increase up to day 4 of adult age, and then incrementally decrease to day 8. However, the magnitude of increase was contingent on the mtDNA haplotype, with only two haplotypes exhibiting a clear peak in reproductive success at day 4 (MYS and ORE). The reaction norms per haplotype crossed-over across the eight days of the experiment, with several haplotypes that exhibited the highest relative reproductive success at the peak of the assay (day 4) generally associated with low reproductive success relative to the other haplotypes at Day 1 and 8 of the experiment (Figure 2A).

**Figure 1:**
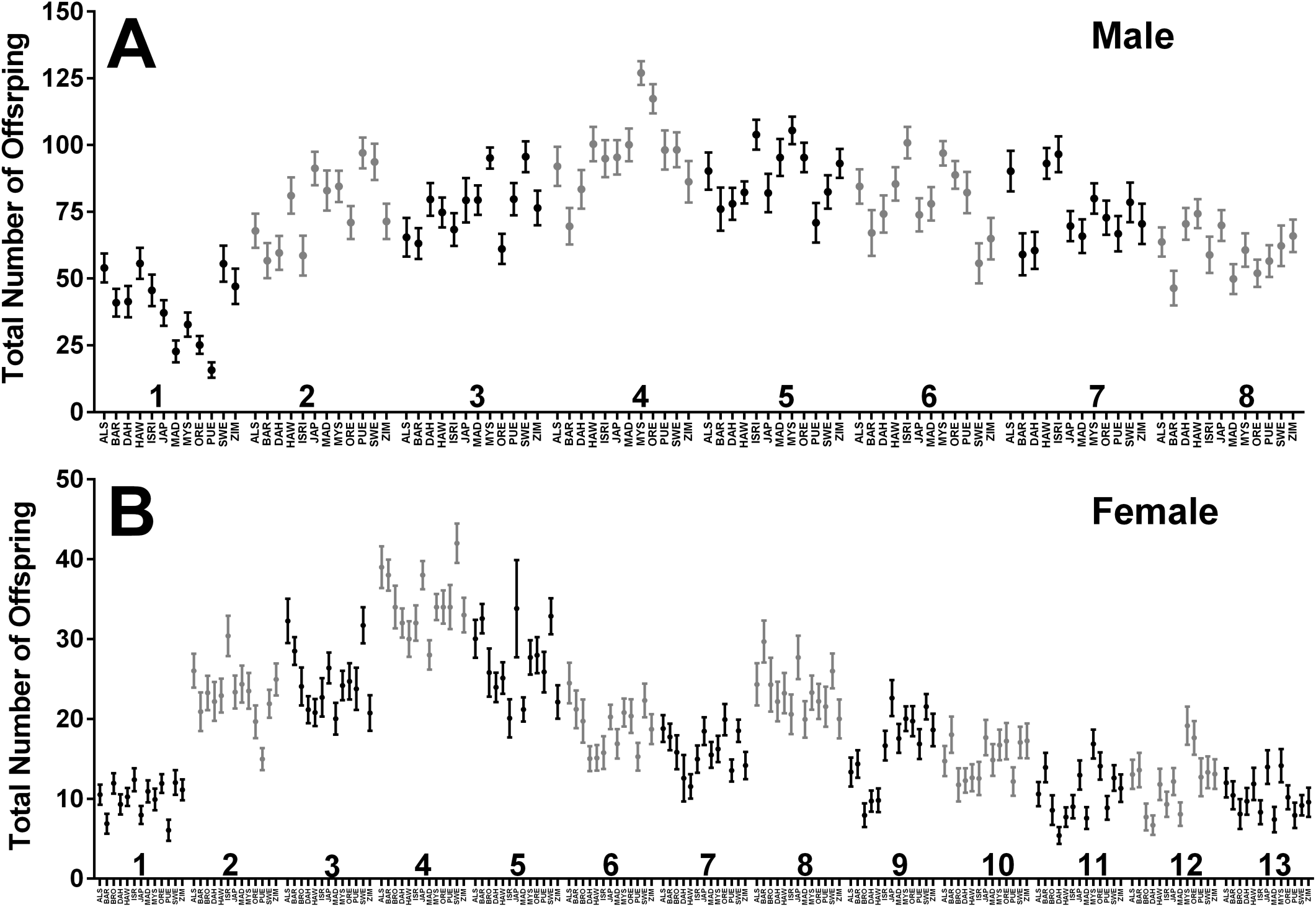
Total number of offspring produced per day (mean ± 1 S.E.) for (**A**) males and (**B**) females across the mitochondrial strains per day of the “sustained offspring production” experiment. The experiment of male offspring production ran for 8 consecutive days, whilst the experiment of female offspring production ran for 13 days.

**Figure 2:**
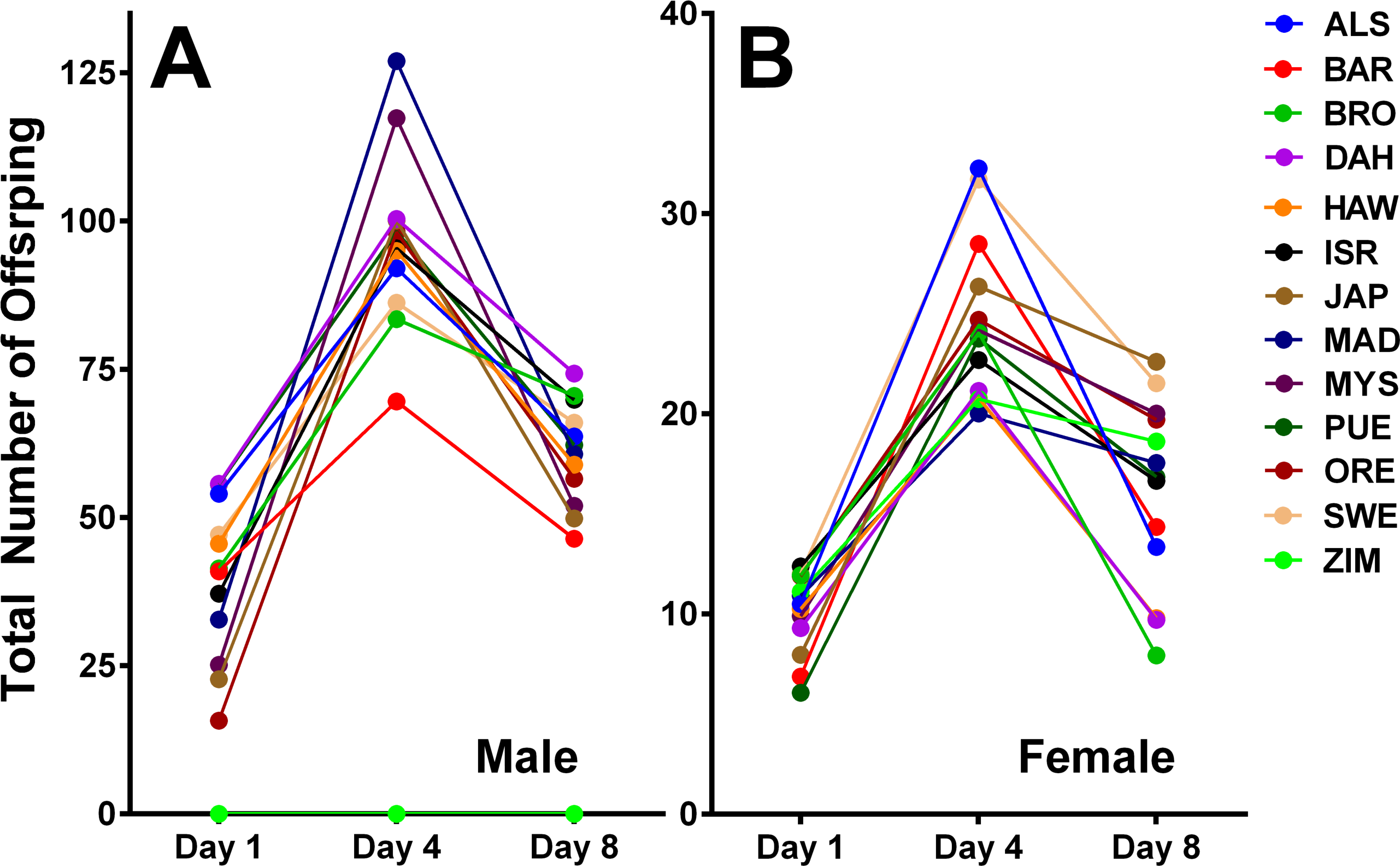
Mean number of offspring produced (reproductive success) for (**A**) males and (**B**) females across the mitochondrial strains, at 3 different age points of the sustained offspring production experiment.

**Table 1:**
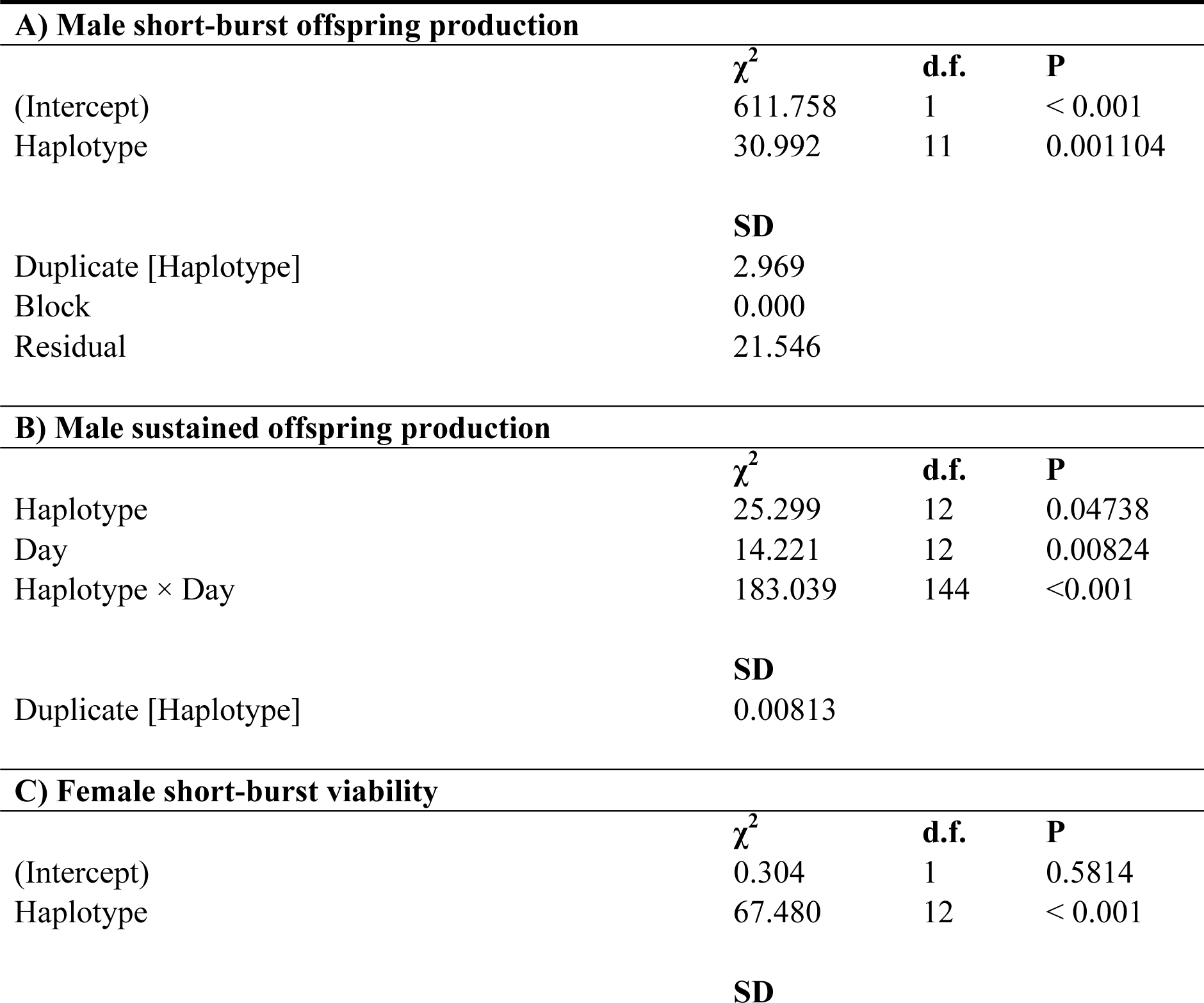
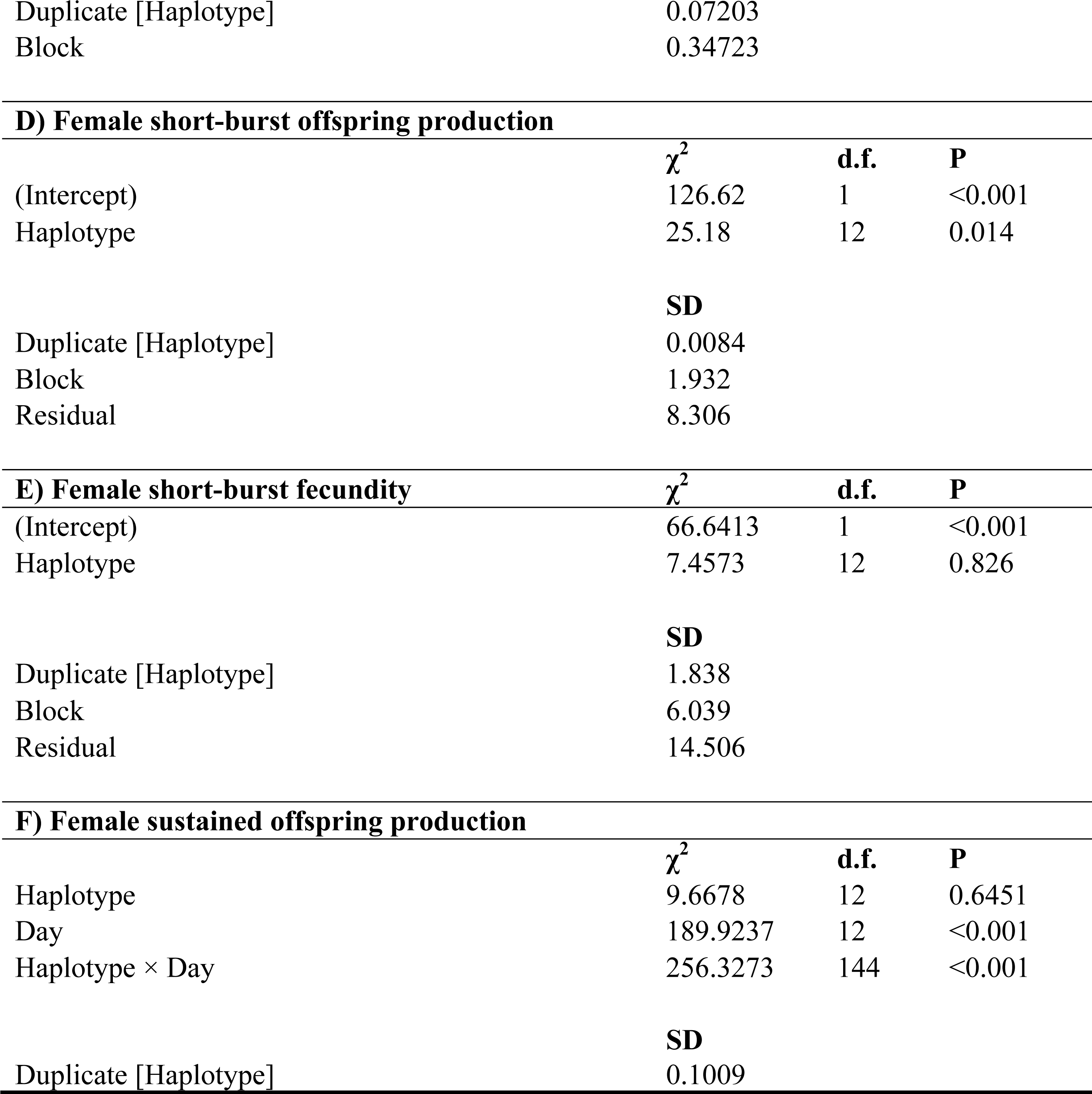
Mitochondrial effects on male (**A**) short-burst offspring production and (**B**) sustained offspring production, and female (**C**) short-burst viability, (**D**) short-burst offspring production, (**E**) short-burst fecundity, and (**F**) sustained offspring production. Haplotype denotes the effect of mitochondrial strain (hence mtDNA haplotype), and Duplicate[Haplotype] denotes the mitochondrial strain duplicate. In the short-burst assays, each experiment was conducted over consecutive sampling blocks (Block). In the sustained offspring production assays, each experiment was conducted over a number of consecutive days (Day; 8 in males, 13 in females). For all models, chi-square test statistics (χ^2^), degrees of freedom, and p values are reported for fixed effects, and standard deviation (SD) for random effects.

| **A) Male short-burst offspring production** |  |  |  |
| --- | --- | --- | --- |
| | $\chi^2$ | d.f. | P |
| (Intercept) | 611.758 | 1 | < 0.001 |
| Haplotype | 30.992 | 11 | 0.001104 |
| **SD** |  |  |  |
| Duplicate [Haplotype] | 2.969 |  |  |
| Block | 0.000 |  |  |
| Residual | 21.546 |  |  |
| **B) Male sustained offspring production** |  |  |  |
| | $\chi^2$ | d.f. | P |
| Haplotype | 25.299 | 12 | 0.04738 |
| Day | 14.221 | 12 | 0.00824 |
| Haplotype × Day | 183.039 | 144 | <0.001 |
| **SD** |  |  |  |
| Duplicate [Haplotype] | 0.00813 |  |  |
| **C) Female short-burst viability** |  |  |  |
| | $\chi^2$ | d.f. | P |
| (Intercept) | 0.304 | 1 | 0.5814 |
| Haplotype | 67.480 | 12 | < 0.001 |
| **SD** |  |  |  |

|  |  |
| --- | --- |
| Duplicate [Haplotype] | 0.07203 |
| Block | 0.34723 |

**D) Female short-burst offspring production**
| | $\chi^2$ | d.f. | P |
| --- | --- | --- | --- |
| (Intercept) | 126.62 | 1 | <0.001 |
| Haplotype | 25.18 | 12 | 0.014 |

**SD**
|  |  |
| --- | --- |
| Duplicate [Haplotype] | 0.0084 |
| Block | 1.932 |
| Residual | 8.306 |

**E) Female short-burst fecundity**
| | $\chi^2$ | d.f. | P |
| --- | --- | --- | --- |
| (Intercept) | 66.6413 | 1 | <0.001 |
| Haplotype | 7.4573 | 12 | 0.826 |

**SD**
|  |  |
| --- | --- |
| Duplicate [Haplotype] | 1.838 |
| Block | 6.039 |
| Residual | 14.506 |

**F) Female sustained offspring production**
| | $\chi^2$ | d.f. | P |
| --- | --- | --- | --- |
| Haplotype | 9.6678 | 12 | 0.6451 |
| Day | 189.9237 | 12 | <0.001 |
| Haplotype × Day | 256.3273 | 144 | <0.001 |

**SD**
|  |  |
| --- | --- |
| Duplicate [Haplotype] | 0.1009 |

### Female Mitochondrial Reproductive Success, and Short-burst Viability Assays

Polymorphisms within the mitochondrial genome affected egg-to-adult viability of a female's clutch (χ^2^ = 67.480, p < 0.001, Table 1C), short-burst offspring production (χ^2^ = 25.18, p = 0.014, Table 1D), but not short-burst fecundity (χ^2^ = 7.4573, p = 0.826, Table 1E). An interaction between mitochondrial strain and day of the mating assay affected sustained female reproductive success (haplotype × day, χ^2^ = 256.3, p<0.001, Table 1F, Figure 1B). All haplotypes exhibited a similar trend, with reproductive success incrementally increasing up until day 4 of the assay, following which point, reproductive success began to decline. Again, however, these patterns were contingent on the mtDNA haplotype, with norms of reaction crossing per haplotype across Days 1, 4 and 8 of the assay (Figure 2B).

### Mitochondrial Genetic Correlations

Intra-sexual correlations between reproductive traits tended to be strongly positive in direction (e.g. r_female short-burst offspring production vs female sustained_ = 0.55; r_male short-burst vs male sustained_ = 0.62, Figure 3). Furthermore, short-burst viability exhibited a strong positive correlation with short-burst offspring production in females, across haplotypes (Figure 3). In contrast, inter-sexual correlations tended to be negative in direction. In particular, the correlations between female and male short-burst offspring production, and between female short-burst viability and male short-burst offspring production, were strongly negative (r_female short-burst offspring viability vs male short-burst offspring production_ = −0.43, Figure 3), as was the inter-sexual correlation between female short-burst fecundity and male sustained offspring production (r_female short-burst fecundity vs male sustained_ = −0.44, Figure 3).

**Figure 3:**
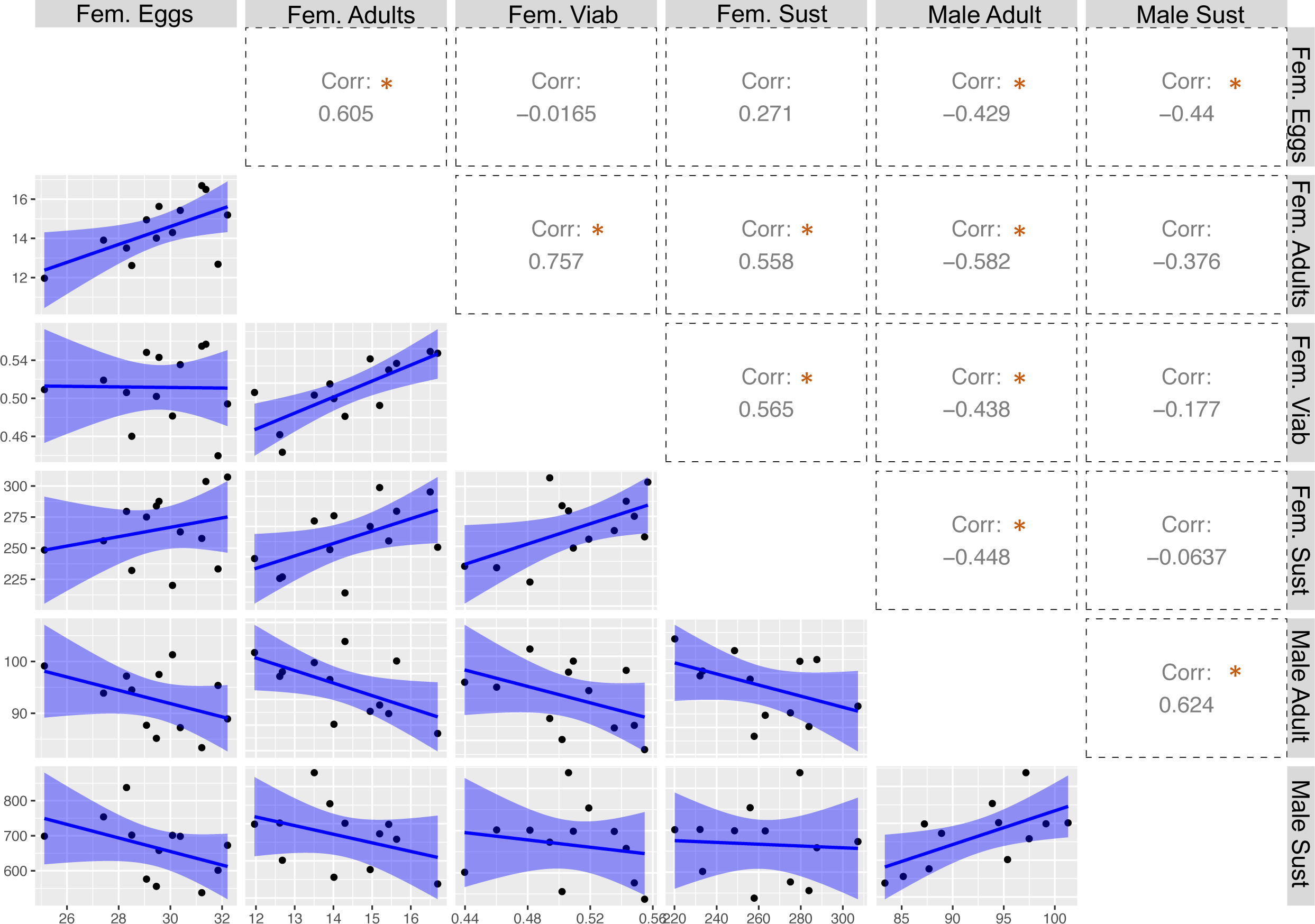
Correlation estimates of intra- and inter-sexual genetic correlations for male and female reproductive traits across mitochondrial haplotypes. Significance is noted as a star next to the coefficient, and represent bootstrapped confidence intervals (95%) that do not overlap with zero.

## Discussion

We explored mitochondrial genetic effects, across distinct and naturally-occurring mitochondrial haplotypes, on components of reproductive success in male and female *D. melanogaster*, using an approach that enabled us to unambiguously trace the genetic effects to the level of the mtDNA sequence. Notably, genetic polymorphisms located across these haplotypes affected almost all components of reproductive success measured – in females and in males. Furthermore, we uncovered strong pleiotropy in the reported effects. These patterns of pleiotropy were positive for intra-sexual correlations across haplotypes (e.g. for associations between short-burst and sustained components of reproductive success in each of the sexes), but negative for several of the inter-sexual correlations.

Negative inter-sexual correlations are striking because they indicate that, at the level of whole haplotypes, those haplotypes that confer relatively high reproductive success in one sex, generally confer low success in the other. Furthermore, we note that our estimate of this negative correlation is conservative, because it excluded the Brownsville mtDNA haplotype, which is completely male-sterile in the nuclear background assayed here (*w^1118^*), and which we have previously reported to host a sexually antagonistic polymorphism located in the *CytB* gene (21). The negative correlation between male and female reproductive success is consistent with evolutionary theory first developed by Frank and Hurst (1996), and which is routinely called “Mother's Curse” (29), which proposes that maternal inheritance of the mitochondria will lead to the accumulation of male-biased mutation loads within the mtDNA sequence (31). Specifically, however, while Frank and Hurst (1996) envisaged that such mutations would accumulate under mutation-selection balance (i.e. the mutations would be largely benign, or slightly deleterious, in their effects on females), our results suggest a role for sexually antagonistic selection (30, 59), with mutations accumulating in the mtDNA sequence that augment female reproductive success, but that come at cost to male reproductive performance.

Under strict maternal inheritance, female-harming but male-benefiting mtDNA mutations should be efficiently purged by purifying selection. In contrast, if mtDNA mutations appear that are female-benefiting, but male-harming, they will presumably increase in frequency under positive selection (30). Furthermore, the pool of sexually antagonistic mutations accumulating within the mitochondrial genomes will differ across populations – in terms of the identity of the mutation sites at which they occur, the associated nucleotides, and total number of mutations accrued. Consequently, at the level of whole haplotypes sourced from different global populations, we should then expect to observe a negative genetic correlation, with haplotypes that harbour numerous female-benefiting but male-harming mutations (or alternatively harbouring a few mtDNA mutations of major sexually antagonistic effect) conferring higher relative female, but lower male, reproductive success. Conversely, those haplotypes harbouring few such mutations (or alternatively mutations of only minor effect) will confer lower female reproductive success relative to other haplotypes, but relatively higher success in males.

In our study, we included egg-to-adult viability of the female clutch, as a measure in our analyses; a measure that lies at the interface between a maternal and an offspring trait (48-53). We found that mitochondrial haplotypic covariance between short-burst viability and female sustained offspring production was strongly positive, while mitochondrial covariance between short-burst viability and male short-burst offspring production was strongly negative. These patterns of covariation are striking, because they indicate that the direction of selection on mitochondrial mutations might not only be directly antagonistic between adult males and adult females, but also between juvenile components of fitness and components of adult male fitness, thus acting to exacerbate the rate at which male-biased mitochondrial mutation loads will accumulate within populations. That is, these correlations imply that mutations exist within the mitochondrial genome that not only augment adult female reproductive success, but also the survival chances of females during juvenile development, and that these mutations will thus presumably be under strong positive selection, even though these same mutations appear to exert negative effects on male components of reproductive success.

Our findings provide strong empirical evidence for the emerging realization that non-neutral polymorphisms that accumulate within the mitochondrial genome will exert pleiotropic effects across life-history traits, not just within a sex, but also across the sexes. Here, in this study, we limited our investigation to correlations across different components of reproductive success in each of the sexes, and also the short-burst viability of reproducing females. But, previous studies have reported mitochondrial genetic associations between longevity and reproductive success, or traits associated with juvenile components of fitness (32, 37, 60). For example, consistent with the signature of intra-sexual positive pleiotropy identified in our experiment here, Dowling et al. (2009) reported a positive genetic association between female longevity and female reproductive success, between two mtDNA haplotypes that were segregating within a population of *D. melanogaster*. Moreover, consistent with the intersexual negative correlations found in our current study, Rand et al. (2001) reported a negative correlation between the sexes for a measure of juvenile viability in *D. melanogaster* (based on a chromosome segregation assay), across two of three mtDNA haplotypes measured. And recently, Camus et al (2015) identified sex-specific effects tied to polymorphisms at key protein-coding mtDNA genes, *mt:ND5* and *mt:Cyt-b*. In particular, a single nonsynonymous mutation (Ala-278-Thr) in the *mt:Cyt-b* gene was associated with patterns of antagonistic pleiotropy both within and between the sexes (21). Females with the Brownsville haplotype, which carries this SNP, were fully fertile but suffered short longevity relative to females with other haplotypes. Males with this SNP, however, suffer reduced fertility (38), and are in fact completely sterile when this SNP is placed in the *w^1118^* nuclear background used here, but experience higher longevity than males with other haplotypes (21). This is the first identified SNP within the mitochondrial genome associated with overtly sexually antagonistic effects. However, as our results here suggest, there are likely to be numerous other such SNPs segregating within the mitochondrial genome.

Finally, we note that the aforementioned Ala-278-Thr SNP in the *mt:Cyt-b* is an example of a SNP in the mtDNA that is putatively entwined in the regulation of the classic life-history trade-off between investment into reproduction versus survival. Our current study reinforces a growing body of evidence that supports the contention that the genetic variation harboured within the mitochondrial genome is likely to influence patterns of covariation between numerous components of life-history in each of the sexes (21, 37, 60). We encourage future studies of mitochondrial genetic variation to increasingly take a multivariate approach when screening for mtDNA-mediated effects on each of the sexes, to help elucidate the extent to which the mitochondrial genome contributes to the evolution of sex differences and life-history trade-offs (32, 61).

## Acknowledgements

We thank Winston Yee for his help with fly husbandry, and David Clancy for providing *Drosophila melanogaster* mitochondrial populations in 2007. The study was funded by the Australian Research Council (grant number DP1092897 and DP170100165 to DKD).

